# Three-dimensional vascular microenvironments uncover endothelial plasticity during TGF-β2-driven vascular remodeling

**DOI:** 10.64898/2026.08.04.742924

**Authors:** Yuning Fu, Kao Tsuchiya, Yuji Nashimoto, Kazuki Takahashi, Yujin Ohsugi, Sayaka Katagiri, Takeshi Hori, Miho Kobayashi, Shotaro Yoshida, Fumiko Itoh, Tetsuro Watabe, Hirokazu Kaji

**Affiliations:** Laboratory for Biomaterials and Bioengineering, Institute of Science Tokyo, Tokyo, Japan; Graduate School of Science and Engineering, Chuo University, Tokyo, Japan; Graduate School of Medical and Dental Sciences, Institute of Science Tokyo, Tokyo, Japan; Institute of Liberal Arts and Science, University of Toyama, Toyama, Japan; Research Center for Autonomous Systems Materialogy (ASMat), Institute of Integrated Research (IIR), Institute of Science Tokyo, Tokyo, Japan; Division of Genome Safety Science, Center for Biological Safety and Research (CBSR), National Institute of Health Sciences (NIHS), Kanagawa, Japan; Laboratory of Stem Cells Regulation, Tokyo University of Pharmacy and Life Sciences; Section of Oral-Systemic Health, Oral Science Center, Institute of Science Tokyo, Tokyo, Japan

**Keywords:** Tumor microenvironment, TGF-β, Vascular network, Microphysiological system, Organ-on-a-chip, Microfluidic device

## Abstract

The tumor microenvironment plays a pivotal role in tumor development, harboring elements such as endothelial cells, immune cells, fibroblasts, and soluble factors such as transforming growth factor-β (TGF-β) family. TGF-β family regulates cell development and promotes tumor invasion, metastasis, angiogenesis, and endothelial-to-mesenchymal transition (EndoMT). Here, we investigate the effects of TGF-β signaling on vascular remodeling using a three-dimensional (3D) vascular network in a microfluidic device. Using both a co-culture (3D-Co) and simplified endothelial monoculture (3D-CM), we demonstrate that TGF-β signaling reduces the quality and functionality of the vasculature by regressing them. In addition, we observed the upregulation of EndoMT-related markers in mRNA and protein expressions, suggesting the induction of EndoMT in 3D vascular networks. The increased vascular permeability stimulated by TGF-β2 also supports the loss of endothelial identity in the 3D-Co. Transcriptomic analysis revealed the coordinated activation of pathways associated with cell migration and EndoMT, along with the suppression of cell cycle progression. A comparative analysis of two-dimensional (2D) and 3D cultures revealed a fundamentally distinct endothelial response to TGF-β2 in the 3D context, including metabolic reprogramming. These findings demonstrate that the 3D microenvironment critically modulates endothelial responses to TGF-β and enables the emergence of vascular phenotypes not captured in 2D systems. This study provides a more physiologically relevant platform to investigate endothelial dysfunction and vascular remodeling.

## Introduction

The tumor microenvironment (TME) is a dynamic and complex ecosystem comprising immune cells, endothelial cells (ECs), fibroblasts, adipocytes, and noncellular components such as the extracellular matrix (ECM), soluble factors, and exosomes^1^. A defining characteristic of the TME is its abnormal vasculature, which results from disrupted signaling such as increased levels of angiogenic factors^2^.

Endothelial-to-mesenchymal transition (EndoMT) plays a crucial role in the development of abnormal vasculature, as ECs in the TME acquire mesenchymal-like traits under the influence of pro-inflammatory and pro-angiogenic signals^3,4^. EndoMT contributes to vessel leakiness, ECM remodeling, and the formation of fibroblast-like cells, further perpetuating abnormal vascular remodeling and enhancing tumor aggressiveness. Studies using mouse models have confirmed the presence of the EndoMT process in disease development, such as cardiac fibrosis *in vivo*, further suggesting that targeting EndoMT could hold therapeutic potential^5^. However, understanding how EndoMT contributes to the progression of various diseases remains a challenge. To dissect complex tissue ecosystems in TMEs, simple *in vitro* models have been utilized as powerful tools for investigating EndoMT.

Various types of ECs cultured in two-dimensional (2D) dishes can induce EndoMT by transforming growth factor-β (TGF-β), a signal that is involved in many cellular processes through various mediators such as receptors, transcription factors, and microRNAs^6–10^. A recent study on the TGF-β effect in 2D culture has emphasized a transitional EndoMT phase known as partial EndoMT in TGF-β2-induced EndoMT reporter ECs^11^. However, traditional 2D culture systems confine cells to rigid, flat environments that lack the essential elements of physiological blood flow, luminal structure, and the ECM. This has led to limitations in capturing the dynamic process of EndoMT in ECs and the contextual significance of signaling within different tissue origins^12–14^.

Recent advances in microphysiological systems (MPS) have transformed the ability to create miniaturized human tissues with customized three-dimensional (3D) structures, physiological cell interfaces, and finely controlled chemical and physical microenvironments^15,16^. MPS enables the creation of 3D models with controlled shear stress, ECM composition, and interstitial fluid flow to allow real-time observations and accurate replication of tumor-endothelial interactions. Using a pump system to generate specific intravascular flow patterns, oscillatory, or disturbed shear stress on the apical side of ECs has been shown to promote EndoMT in cerebral arteriovenous malformations by activating Notch3 signaling^17,18^. To investigate the critical role of fluid dynamics in the extravascular region, several studies have mimicked the ECM by incorporating various soluble factors, proteins, and exosomes. Mina et al. developed an MPS that integrated an ECM environment to study the combined effects of shear stress and TGF-β on EndoMT^19^. Using the same design, they demonstrated that altered ECM composition, such as variations in collagen levels, glycosaminoglycan concentrations, and matrix stiffness, can independently induce EndoMT, highlighting the importance of the biochemical and mechanical properties of the ECM^20^.

However, the vascular networks used in the study were prepatterned rather than spontaneously formed, presenting challenges in evaluating EndoMT owing to the absence of key spontaneous characteristics inherent to natural human vasculature. The spontaneously formed vasculature closely mimics the natural processes of vascular development, including cell self-organization, dynamic remodeling, and interaction with the surrounding microenvironment^21–23^. While MPS have been widely utilized for studying EndoMT, it remains unclear whether 3D vascular organization alters endothelial responses to TGF-β. Moreover, no studies to date have examined EndoMT using spontaneously formed vascular networks that closely recapitulate vascular self-organization and remodeling.

In this study, we utilized an *in vitro* vascular network fabricated using a five-channel microfluidic device^24^ to co-culture human lung fibroblast cells (hLFs) and human umbilical vein endothelial cells (HUVECs). This co-culture facilitated the spontaneous formation of thick, lumen-like vessels. Using this platform, we systematically investigated the effects of TGF-β2-induced EndoMT under physiologically relevant 3D conditions. We optimized TGF-β2 stimulation and identified concentration-dependent vascular remodeling responses characterized by vascular thinning and increased permeability. HUVECs stimulated by 3 ng/mL of TGF-β2 induced the most pronounced EndoMT-associated phenotypes, as confirmed by both morphological quantification and gene expression profiling. The EndoMT progression was confirmed by gene expression profiles. Higher permeability was observed in 3 ng/mL of TGF-β2-treated vascular network, further confirming the role of TGF-β2 in impairing the endothelial function. To elucidate the underlying mechanisms of 2D and 3D models, we performed a comprehensive gene expression analysis, revealing substantial differences in signaling responses and highlighting the importance of the microenvironmental context in regulating EndoMT. Finally, we validated the physiological relevance of our model by comparing vascular morphological changes with those observed in an *in vivo* model, demonstrating consistent responses to TGF-β2 stimulation. Together, this study provides a robust 3D platform for modeling EndoMT and offers new insights into the regulation of endothelial plasticity in physiologically relevant environments.

## Materials and methods

### Device fabrication

A microfluidic device was employed to prepare a 3D vascular network, similar to that used in previous reports^24^. Microfluidic devices were fabricated via standard soft lithography. The photomask (TOPIC) was designed using AutoCAD software (Autodesk). SU-8 3000 (Nippon Kayaku) that was 100 µm in height was spin-coated on a silicon wafer (Canosis) and the mold was treated with trichloro (1H,1H,2H, and 2H-perfluorooctyl) silane (Merck).

Polydimethylsiloxane (PDMS, SILPOT 184 W/C, Dow Inc.) and the curing reagent were mixed in a 10:1 weight ratio and poured into the SU-8 mold, followed by degassing and curing overnight at 75 °C, and then peeled off from the mold. Inlets and outlets of each microchannel were prepared using a 2 mm biopsy punch (Kai). The PDMS was bonded to a cover glass (Matsunami Glass) following oxygen plasma treatment (Yamato Scientific). The microfluidic devices were heated overnight at 75 °C and sterilized by UV irradiation for 30 min before use.

### Mice

The control mice (TβRII^fl/fl^) and endothelial cell-specific TGF-β type II receptor-deficient mice (TβRII^iΔEC^) were generated and treated as previously described^25^. The retinal vasculature was selected because its planar architecture enables ready visualization and quantitative analysis of the entire vascular network. The animal procedures were approved by the Institutional Animal Care and Use Committee at Tokyo University of Pharmacy and Life Sciences (approval number: L16-13, LS27-007). Experiments were performed according to the guidelines of the Animal Care Standards of the Tokyo University of Pharmacy and Life Sciences. TβRII^fl/fl^; Pdgfb-iCreER mice were administered tamoxifen at postnatal days 1 and 3 (P1 and P3), and the retinas were harvested and analyzed at P3.

### Cell culture and formation of a 3D vascular network

Green fluorescent protein-labeled human umbilical vein endothelial cells (GFP-HUVECs) were purchased from Angio-Proteomie (cAP-0001GFP) and cultured in endothelial cell growth medium-2 (EGM-2, Lonza) supplemented with 5% penicillin-streptomycin (Thermo Fisher Scientific). Human lung fibroblasts (hLFs) were purchased from Lonza (CC-2512) and cultured in fibroblast growth medium-2 (FGM-2, Lonza) supplemented 5% P/S. Passage numbers 5–6 were used in all experiments for both GFP-HUVECs and hLFs.

To create a 3D co-culture vascular network (3D-Co), GFP-HUVECs and hLFs were seeded into the microfluidic device at 8.0 × 10^6^ and 5.0 × 10^6^ cells/mL, respectively, similar to previous reports^24^. To exclude the effects of hLFs, we used a 3D vascular network formed by hLF-conditioned media (CM) with only GFP-HUVECs. This model is hereafter referred to as the 3D CM-driven vascular network (3D-CM). After a seven-day culture, both models were stimulated by adding TGF-β2 (Thermo Fisher Scientific) (hereafter termed TGF-β) for four days (Figs. 1a and 2a). Detailed information regarding cell seeding into the microfluidic device, 2D cell culture, and TGF-β treatment is provided in Supporting Information.

**Fig. 1.**
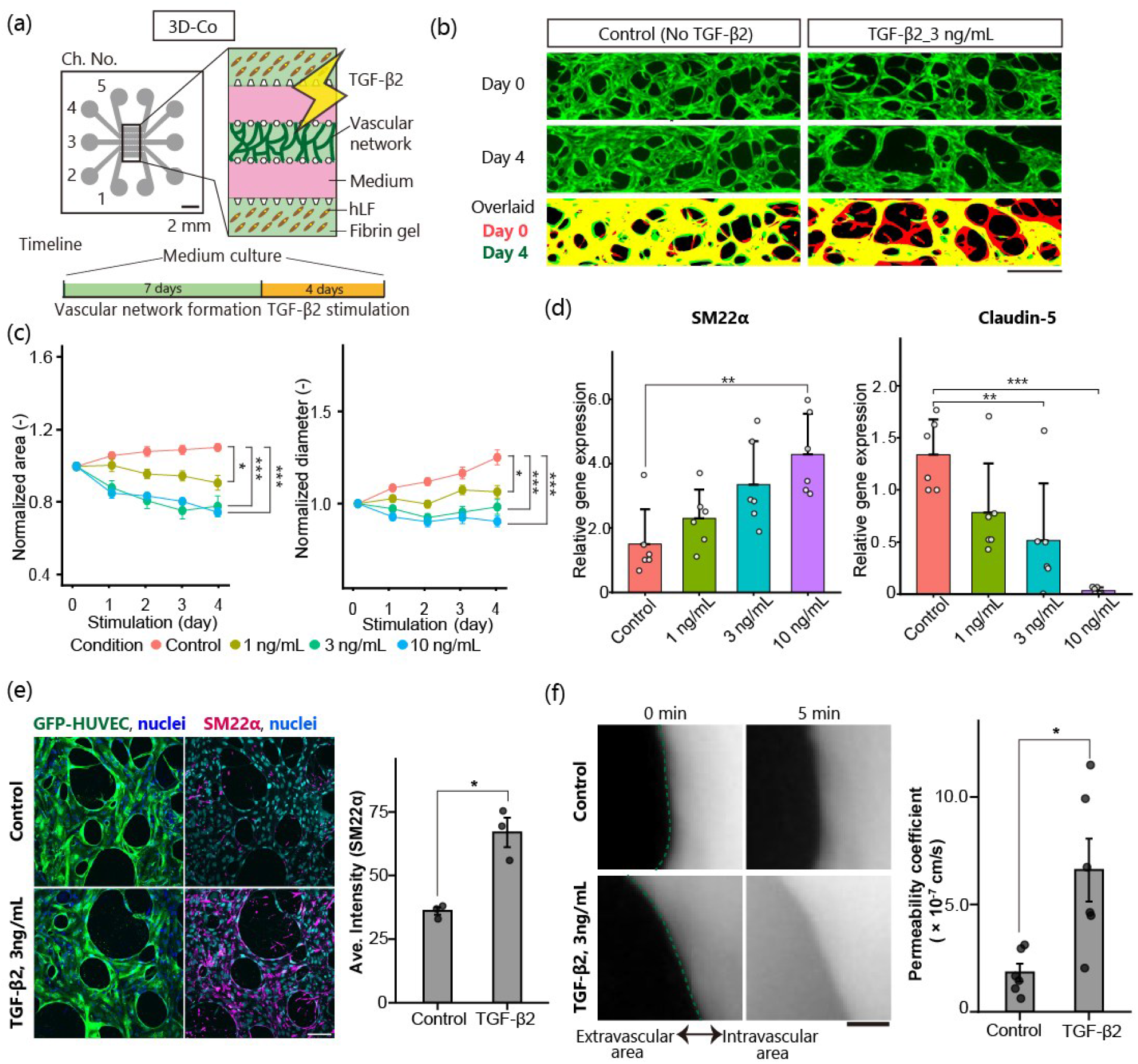
Morphological, molecular, and functional alterations by TGF-β2 in 3D co-culture vascular network (3D-Co). (a) Schematic of cell embedding within the microfluidic device. (b) Representative images showing regression of vascular area following 3 ng/mL TGF-β2 stimulation. Red indicates the binarized vascular area at day 0 (start of stimulation), and green represents the binarized vascular area at day 4 (end of stimulation). Overlapping regions appear yellow. Scale bar: 500 μm. (c) Changes in vascular area (left) and diameter (right) after treatment with TGF-β2. All quantification was normalized to day 0 values. (d) qRT-PCR analysis of the expression of a mesenchymal cell marker SM22α (*TAGLN*), and endothelial cell marker Claudin-5 (*CLDN5*). All data are normalized to the *ACTB* expression, n = 6. (e) Immunofluorescent analysis. Staining for SM22α (magenta) and nuclei (blue) (left). Scale bar: 100 μm. Comparison of the average intensity of SM22α (right), n = 3. TGF-β2 is 3 ng/mL. (f) Vascular permeability assay. Representative fluorescence images (left) display the distribution of TRITC-dextran within the vascular regions. The dashed line demarcates the extravascular (left) and intravascular (right) compartments. Scale bar: 50 µm. The corresponding quantification of the permeability coefficient is presented in the bar plot (right), n = 6. TGF-β2 is 3 ng/mL. All data are shown as mean ± SD Statistical analyses: one-way ANOVA for Figs. 1c and 1d, *t*-test for Figs. 1e and 1f; \**p* < 0.05; \*\**p* < 0.01; \*\*\**p* < 0.001. Comparisons without asterisks are not statistically significant.

**Fig. 2.**
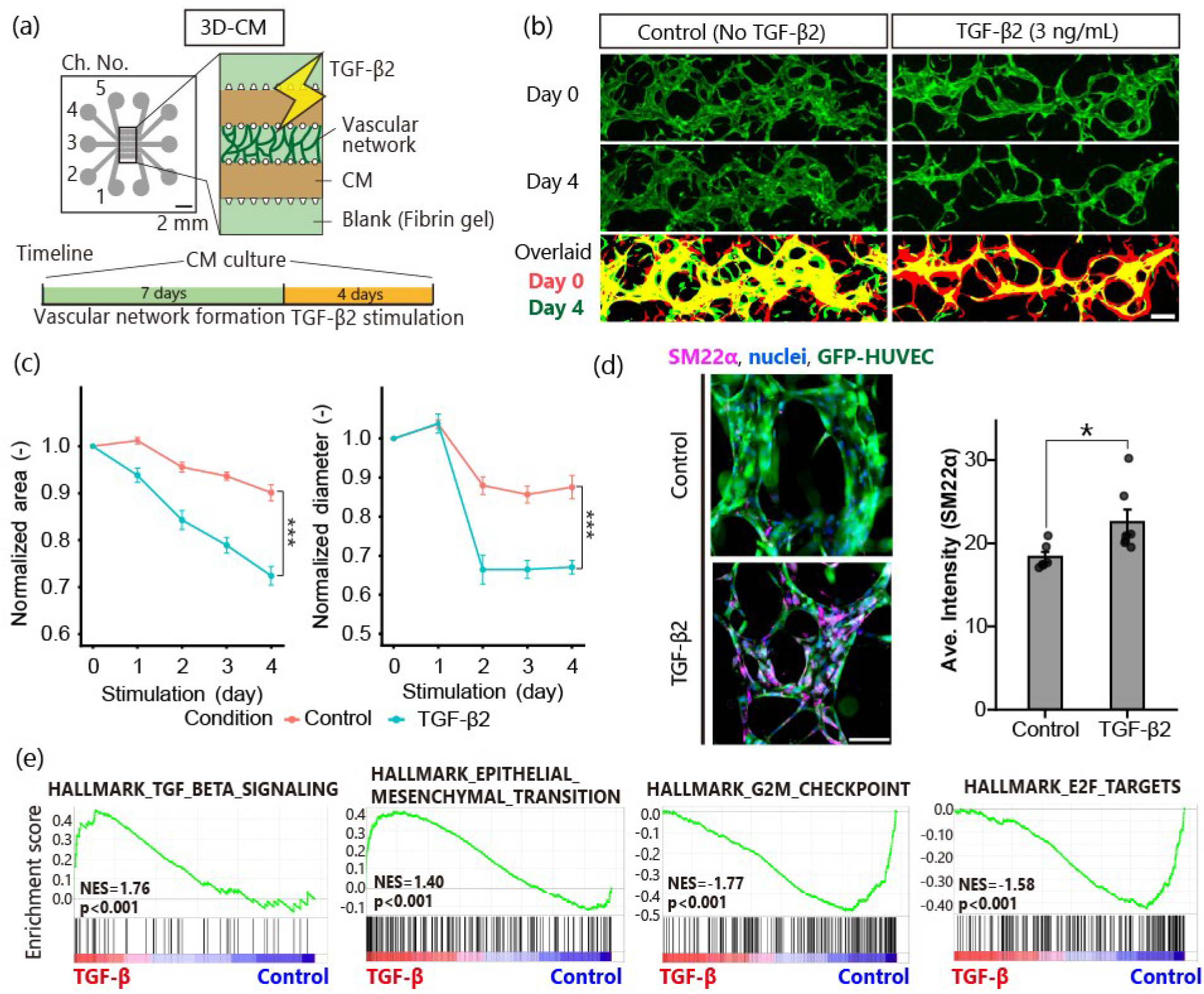
Morphological and molecular alterations by TGF-β2 in 3D CM-driven vascular network (3D-CM). (a) Schematic of cell embedding within the microfluidic device and experiment timeline. CM represents LF conditioned medium. Blank is fibrin gel. (b) Representative images showing a reduction in the vascular area formed by GFP-HUVECs after treatment with 3 ng/mL of TGF-β2 in 3D-CM. Red indicates the binarized vascular area at day 0 (start of stimulation), and green represents the binarized vascular area at day 4 (end of stimulation). Overlapping regions appear yellow. Scale bar: 200 μm. (c) Vascular area (left) and diameter (right) changes from day 0 to day 4. All the quantifications were normalized to the day 0 value, n = 13. Statistical analysis: Welch’s unpaired t-test; ***p < 0.001. (d) Immunocytochemical analysis. Staining for SM22α (magenta) and nuclei (blue) (left). GFP-HUVECs are shown in green. Scale bar: 100 μm. Average intensity of SM22α (right), n = 6. All data are shown as mean ± SD. Statistical analyses: t-test; *p < 0.05; ***p < 0.001. (e) Gene set enrichment analysis of TGF-β signaling, epithelial-mesenchymal transition, and proliferation gene signatures, comparing RNA-seq data from 3D-CM treated without (control) and with TGF-β2. n = 4 for each group. NES, normalized enrichment score; q, FDR-adjusted p-value.

### Cell staining and imaging

The 2D ECs and 3D vascular network were fixed using 4% paraformaldehyde (FUJIFILM Wako Pure Chemical) in phosphate-buffered saline (PBS, Wako) (2D: 30 minutes at room temperature, 3D: 1 h at 4 ℃). Permeabilization was processed with 0.1% Tween 20 (Merck) in PBS (2D: more than 1 h at 4 ℃, 3D: more than 2 h at 4 ℃) and then washed with PBS. Primary antibodies (SM22α (Anti-TAGLN, Abcam, 1:1000), Ki-67 (Abcam, 1:900)) were diluted in 1% bovine serum albumin (BSA) in PBS and added to the dishes and devices. After washing the sample, they were left in PBS for 8 h at 4 ℃. Samples were incubated with 4′,6-diamidino-2-phenylindole, Alexa Fluor 555 donkey anti-rabbit IgG (BioLegend), or Alexa Fluor 647 donkey anti-rabbit IgG (H+L) (Thermo Fisher Scientific) diluted in 1% BSA/PBS for 8 h at 4 °C. After incubation, the samples were washed and immersed in PBS.

To evaluate apoptosis within the 3D vascular networks, caspase-3/7 staining was performed using CellEvent™ caspase-3/7 detection reagents (green/red) (Thermo Fisher Scientific). A 100× stock solution was prepared by reconstituting the caspase-3/7 reagent in 100 μL of PBS. This stock was subsequently diluted 1:10 in complete culture medium to obtain a 10× working solution. The working solution was then applied to 3D vascular network devices and incubated for 45 min at 37 °C.

### Analyze of vascular morphology

Analysis of 3D vascular morphology was performed using Fiji^26^. Fluorescence images of GFP-labeled vascular networks were first converted into binary images using a consistent thresholding method. The binarized area was defined as the total vascular area. Subsequently, the binary images were skeletonized using a skeletonize plugin to extract the vascular centerlines. The branch number and average branch length were quantified using the Analyze Skeleton plugin, and the total vessel length was calculated as the sum of all skeletonized branches. The average vascular diameter was estimated by dividing the total vascular area by the total vessel length as follows:

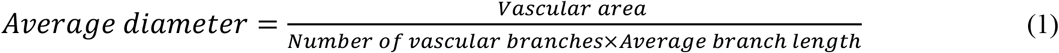

The *in vivo* mice binarized images were processed and analyzed using Fiji and Python. CD31-positive areas were identified as vascular networks. Regions of interest for measuring the vascular diameter and length were manually defined, and quantitative analysis was performed using Fiji.

### qRT-PCR assays

Only the Ch. 3 section of the device was cut out with a scalpel. The vascular network attached to the glass surface was collected with a 1 mm biopsy punch and dissolved in a tube containing 45 μL of Buffer RLT Plus (QIAGEN) with β–mercaptoethanol (Sigma-Aldrich). A total of 7.5 μL of Buffer RLT Plus was applied to both the glass surface and cut-out PDMS surface, and the lysate was immediately collected into a tube. This process was repeated two or three times.

Total RNAs were purified with RNeasy Plus Micro kit (QIAGEN) and converted into cDNA using the PrimeScript II 1st Strand cDNA Synthesis Kit (Takara Bio). Quantitative reverse transcription-polymerase chain reaction (qRT-PCR) analysis was performed using the ABI StepOnePlus system (Applied Biosystems) with PowerUp™ SYBR™ Green Master Mix (Thermo Fisher Scientific). All expression data were normalized to β-actin expression. Primer sequences are listed in Supplementary Table 1. All primers were synthesized by Eurofins Genomics.

### RNA-seq analysis

Total RNAs from HUVECs cultured in 2D dishes and 3D-CM were collected at the end of TGF-β stimulation using the RNeasy Plus Micro Kit (QIAGEN). The RNA sequencing (RNA-seq) was outsourced to Rhelixa Corp. RNA-seq libraries from microfluidic devices were prepared from 2D cultured ECs using SMART-Seq mRNA HT LP (Takara) and NEBNext Ultra II Directional RNA Library Prep Kit (New England Biolabs) with NEBNext Poly(A) mRNA Magnetic Isolation Module (New England Biolabs). The adjusted libraries were sequenced with 150-bp paired-end reads on a NovaSeq X Plus (Illumina). Low-quality and adapter sequences from paired-end reads were trimmed. The raw reads were mapped to Homo sapiens. Differential expression analysis was conducted using the DESeq2 v1.32.0 package for R. Differentially expressed genes (DEGs) were defined using the following criteria: |fold change (FC)| > 1.5, adjusted p-value < 0.05 and log_2_TPM > 2 for Venn diagram analysis comparing 3D and 2D cultures. The DEGs identified from the Venn analysis were subjected to Gene Ontology (GO) enrichment analysis. Gene Set Enrichment Analysis (GSEA) was performed using the entire ranked gene list, and principal component analysis (PCA) was performed using normalized gene expression data in R.

### Permeability assay

The measurement method is based on previous research^27,28^. Tetramethylrhodamine (TRITC-dextran) (70 kDa, Sigma) was prepared with PBS. The medium in the reservoirs of Chs. 2 and 4 was removed. A total of 10 μL of TRITC-dextran was introduced into Ch. 2, and images were sequentially taken and stored every minute for up to 5 min. The permeability coefficient P (cm/s) was calculated by the following equation:

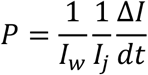

where *I_w_* is the length of the blood vessel to be measured (cm), *I_j_* is the average intensity in the blood vessel, *I* is the total difference of the intensities of the extravascular gel regions in the measurement interval (cm^2^), and *t* is the measurement interval (s).

### Statistical analysis

Values are presented as mean ± standard deviation. Significant differences between means were determined using an unpaired two-tailed Student *t*-test and Mann–Whitney *U* test, as appropriate. Comparisons among multiple groups were performed using one-way analysis of variance followed by Tukey’s post-hoc multiple comparisons test. Differences between means were considered statistically significant at \**p* < 0.05.

## Results

### Morphological and functional changes of 3D co-culture vascular network by TGF-β2 stimulation

To establish a robust 3D vascular model, GFP-HUVECs were co-cultured with hLFs, which provide essential growth factors for vascular formation (3D-Co). TGF-β2 (hereafter termed TGF-β), a member of the TGF-β family, has been identified as the most potent inducer of EndoMT in 2D EC models, with a concentration of 1 ng/mL as particularly effective^29^. Based on these results, a concentration range of approximately 1 ng/mL was tested. Following a seven-day maturation period, the established networks were exposed to varying concentrations of TGF-β (0, 1, 3, and 10 ng/mL) for four days (Fig. 1a). The GFP and binarized overlaid channel images demonstrated that the control group maintained a stable and well-organized morphology characterized by a larger vascular area compared to the TGF-β-treated (3 ng/mL) group (Fig. 1b). Quantitative analysis revealed that the network began to destabilize at a concentration of 1 ng/mL. This effect was markedly amplified at 3 ng/mL, which manifested as the loss of vascular area and a significant reduction in vascular diameter (Figs. 1b and 1c). The inhibitory effects of 10 ng/mL were comparable to those of 3 ng/mL. Given that 10 ng/mL represents a supraphysiological concentration, we identified 3 ng/mL as the optimal concentration for inducing vascular remodeling in this 3D-Co platform.

To examine the molecular mechanism in the response to TGF-β stimulation in 3D-Co, we quantified the expression of endothelial and mesenchymal markers using qRT-PCR. The analysis revealed that TGF-β stimulation increased SM22α (*TAGLN*), the essential marker of mesenchymal cells, and decreased Claudin-5 (*CLDN5*), the endothelial cell marker (Fig. 1d). Other main mesenchymal markers, such as *SNAI1*, *COL1A1*, and *FN1* expression levels, also increased in response to TGF-β stimulation (Supplementary Fig. 1). The induction of *SNAI1*, a core transcription factor typically activated via the Smad2/3 signaling pathway, further supports the activation of the canonical TGF-β signaling axis in our 3D-Co platform. Immunostaining validated these results, showing a significant accumulation of SM22α protein throughout the 3D-Co (Fig. 1e). Collectively, these findings confirm that TGF-β promotes EndoMT, thereby driving the observed structural regression of the 3D vascular network.

Beyond molecular and morphological remodeling, given that EndoMT increases endothelial cell permeability by reducing cell-cell junction proteins^30^, we further evaluated the functional integrity of 3D-Co by assessing its barrier function. In the unstimulated control group, the vasculature exhibited robust barrier properties with minimal leakage of TRITC-dextran (70 kDa). In contrast, TGF-β stimulation triggered a pronounced increase in TRITC-dextran permeation into the extravascular area from 0 to 5 min (Fig. 1f, the permeability coefficient was (1.8 ± 1.0) × 10^−7^ cm/s in the control group, (6.6 ± 1.0) × 10^−7^ cm/s following TGF-β treatment). This nearly fourfold increase in permeability aligns with the observed downregulation of the junctional protein Claudin-5 (*CLDN5*) (Fig. 1d), confirming that TGF-β-induced EndoMT leads to a profound loss of vascular barrier function. Taken together, these results demonstrate that TGF-β drives structural regression and severely impairs the physiological gating capacity and barrier integrity of 3D-Co.

### Characterization of a simplified 3D CM-driven vascular network (3D-CM) for endothelial RNA extraction

Given that 3 ng/mL of TGF-β induced the most significant remodeling in 3D-Co, we aimed to analyze the underlying molecular changes via RNA sequencing. Although sample collection was carefully performed to minimize contamination by hLF-derived RNA, hLFs in Chs. 1 and 5 of 3D-Co tended to migrate into the vascular channel (Ch. 3), posing a risk of contamination of endothelial-specific RNA samples over time. To circumvent this, a 3D-CM model was developed. In this configuration, Chs. 1 and 5 contained a blank fibrin gel, while GFP-HUVECs were seeded in Ch. 3. To simulate the stromal environment, Ch. 2 and 4 were supplied with LF CM instead of live hLFs (Fig. 2a).

To verify the functional fidelity of this simplified model, we compared its response to 3 ng/mL of TGF-β against a control group. Time-lapse images showed a significant decrease in the vascular area in the TGF-β-treated (3 ng/mL) group compared with the control group (Fig. 2b). Quantification revealed a significant reduction in both the vascular area and diameter, mirroring the trends observed in 3D-Co (Figs. 1c and 2c). To investigate whether these changes were driven by the EndoMT, we performed immunofluorescence staining for SM22α. The results showed a marked increase in SM22α expression upon TGF-β stimulation (Figs. 2c and 2d), confirming that the 3D-CM model effectively recaptures both the morphological regression and molecular signaling characteristics of TGF-β-induced vascular remodeling.

To elucidate the molecular basis of the observed vascular regression, we performed transcriptomic analysis on the vascular network with or without TGF-β stimulation (3 ng/mL) in the 3D-CM model. GSEA revealed a marked enrichment of the TGF-β signaling hallmark, confirming the activation of TGF-β signaling in response to TGF-β treatment. The epithelial-mesenchymal transition (EMT) hallmark, resembling upregulated genes in EndoMT-induced cells was also significantly enriched (Fig. 2e). GO enrichment analysis demonstrated the upregulation of pathways associated with cell migration (Supplementary Fig. 2a). Together, these transcriptional changes suggest that ECs acquire a motile, mesenchymal-like phenotype, which is indicative of EndoMT in the 3D-CM model. In parallel, GSEA indicated a downregulation of cell cycle-related gene sets (G2M CHECKPOINT, Fig. 2e), suggesting the suppression of cell cycle progression upon TGF-β stimulation. However, immunostaining for the proliferation marker Ki-67 did not reveal a statistically significant difference between groups (Supplementary Figs. 2b and 2c), indicating that the transcriptional changes might not fully translate into detectable alterations at the protein or cellular level under the current experimental conditions.

GO analysis identified a significant enrichment of pathways related to programmed cell death and apoptosis (Supplementary Fig. 2a). To validate this observation, we performed fluorescence staining of caspase-3/7. The results showed a significant increase in caspase-3/7-positive cells in TGF-β-treated vessels compared with controls (Supplementary Figs. 2b and 2c), supporting the induction of endothelial apoptosis.

Taken together, these findings suggest that TGF-β promotes vascular regression in the 3D-CM model through two coordinated mechanisms. First, it induces EndoMT, as evidenced by transcriptional signatures of EMT and enhanced migratory pathways, leading to alterations in endothelial identity. Second, TGF-β promotes apoptosis and suppresses proliferative capacity, contributing to the structural destabilization and reduction of vascular networks.

### Comparative analysis between 2D and 3D culture models

To investigate the transcriptomic divergence between the 2D traditional culture and 3D culture environment, we compared the gene expression profiles of 2D sparse (low cell density), 2D confluent (high cell density), and 3D-CM models (Figs. 3a and 3b). Upon stimulation with 3 ng/mL TGF-β, SM22α expression significantly increased in the 2D sparse group, indicating activation of TGF-β signaling (Supplementary Figs. 3a and 3b). In contrast, no significant change was observed in the 2D confluent group. These results suggest that high cell density in confluent cultures may provide protective intercellular signaling that attenuates TGF-β-induced EndoMT.

**Fig. 3.**
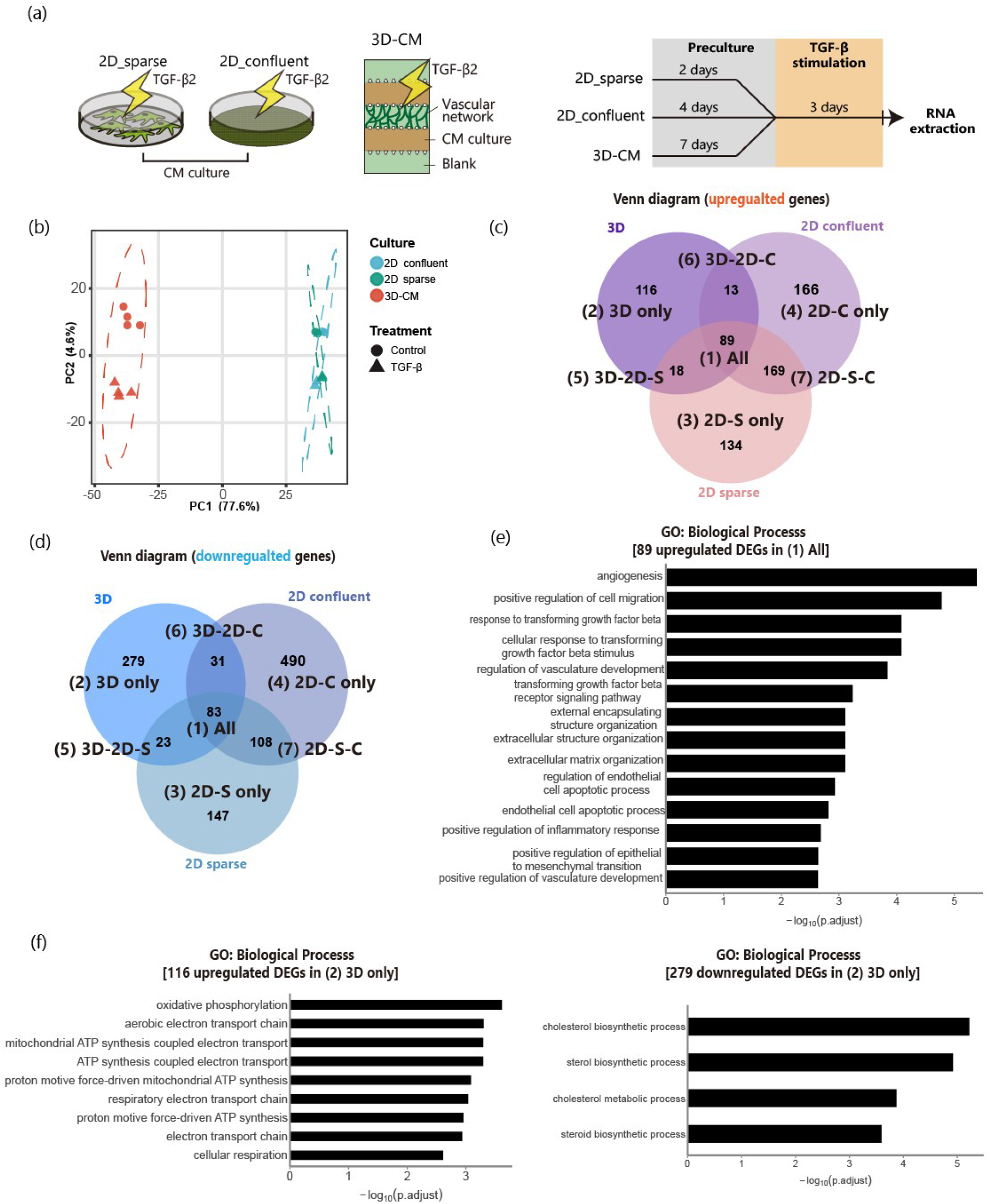
Comparative RNA-seq analysis between 2D sparse, 2D confluent, and 3D-CM culture models. (a) RNA extraction from 2D sparse, 2D confluent, and 3D-CM models. n = 2 for each group in 2D samples, n = 4 for each group in 3D-CM. (b) Principal component analysis (PCA) of transcriptomic profiles. PCA plot showing the global gene expression patterns of human umbilical vein endothelial cells (HUVECs) cultured in 2D (sparse and confluent) and 3D-CM. Each point represents an independent biological replicate. The x- and y-axis represent the first two principal components (PC1 and PC2), accounting for 77.6% and 4.6% of the total variance, respectively. Samples are color-coded by culture condition (green: 2D sparse; blue: 2D confluent; red: 3D-CM). (c) Venn diagram showing number of differentially expressed genes (DEGs) upregulated by TGF-β in 3D-CM, 2D confluent, and sparse HUVECs. (d) Venn diagram showing number of DEGs downregulated by TGF-β in 3D-CM, 2D confluent, and 2D sparse HUVECs. All DEGs input into the Venn diagram were filtered with a stringent threshold of |FC| > 1.5, p-adjust < 0.05, and a base expression level of log_2_TPM > 2 to ensure biological relevance. (e) Gene ontology (GO) functional enrichment analysis of DEGs upregulated by TGF-β in (1), “all”. (f) GO functional enrichment analysis of DEGs upregulated (left) and downregulated (right) by TGF-β in (2), “3D only”. Bar plot showing the significantly enriched GO terms for DEGs under TGF-β regulation. DEGs were identified using the criteria of |FC| > 1.5 and padj < 0.05.

PCA revealed that the first two principal components (PC1 and PC2) accounted for 77.6% and 4.6% of the total variance, respectively (Fig. 3b). The results showed that 3D-CM clustering entirely apart from the 2D culture samples, which indicated PC1 clearly separated the samples by culture dimensionality. However, 2D sparse and 2D confluent samples clustered closely together along the PC1 axis, indicating that increasing the cell density to confluence in 2D was insufficient to bridge the massive transcriptomic shift induced by a 3D environment. The distinct separation between the control and TGF-β-treated samples across all models indicated that PC2 captured the cellular response to TGF-β signaling. Even under nonstimulated conditions, 3D-CM remained clearly separated from both 2D cultures, indicating that the endothelial transcriptome was fundamentally reshaped by the 3D microenvironment before TGF-β stimulation. These findings demonstrate a pronounced transcriptomic divergence between 2D and 3D-CM cultures, highlighting the dominant impact of the 3D microenvironment on endothelial gene expression programs.

The distinct baseline transcriptional programs established in 3D-CM and 2D cultures under nonstimulated conditions prompted a further characterization of the intrinsic gene expression differences between 2D and 3D control groups. Venn analysis identified overlapping DEGs shared between the comparisons of 3D control versus 2D sparse control, and 3D control versus 2D confluent control (Supplementary Figs. 4a and b). GO analysis revealed that genes upregulated in 3D-CM were predominantly associated with mitochondrial metabolism, including oxidative phosphorylation and electron transport-related processes (Supplementary Fig. 4c). In contrast, genes commonly downregulated in 3D-CM were enriched in biological processes related to angiogenesis, tissue morphogenesis, endothelial cell migration, TGF-β signaling, and cell proliferation (Supplementary Fig. 4d), indicating that ECs cultured in the 3D microenvironment exhibit a transcriptional profile distinct from conventional 2D cultures even in the absence of stimulation. GSEA demonstrated significant enrichment of the HALLMARK_INFLAMMATORY_RESPONSE in the 3D-CM control group compared with the 2D culture conditions (Supplementary Figs. 4e and g). Conversely, HALLMARK_TGF_BETA_SIGNALING and HALLMARK_G2M_CHECKPOINT were enriched in the 2D control groups relative to 3D-CM (Supplementary Figs. 4f and h), suggesting that ECs maintained under conventional 2D conditions possess a more proliferative and TGF-β-responsive basal transcriptional state. A comparison between the two 2D culture conditions is shown for reference (Supplementary Figs. 4i and j). Because 2D and 3D-CM cultures exhibited distinct basal transcriptional programs prior to stimulation, we investigated whether the endothelial transcriptional response to TGF-β was similarly shaped by the culture microenvironment. To identify transcriptional changes that were uniquely associated with each culture condition following TGF-β stimulation, uniquely expressed DEGs were identified by Venn analysis for upregulated and downregulated genes (Figs. 3c and 3d). GO enrichment analysis was subsequently performed for each subset.

In the shared DEG subset (All, Fig. 3e), TGF-β stimulation was associated with the enrichment of biological processes related to cell migration (positive regulation of cell migration), vascular development (angiogenesis, regulation of vasculature development), extracellular structure organization, and extracellular matrix organization, as well as inflammatory response (positive regulation of inflammatory response). These processes are broadly consistent with previously reported transcriptional features associated with endothelial activation and EndoMT^31–33^.

In contrast, the 3D-specific DEG subset (3D only, Fig. 3d) revealed a distinct enrichment pattern. Downregulated genes were significantly enriched in lipid and cholesterol biosynthetic processes, including sterol and steroid metabolism. Upregulated genes were predominantly associated with mitochondrial energy metabolism, including oxidative phosphorylation and electron transport chain pathways. These results suggest a shift in the metabolic state characterized by reduced lipid biosynthesis and enhanced mitochondrial respiration. Given the essential role of lipid metabolism in maintaining endothelial membrane composition, junctional integrity, and barrier function, these transcriptional changes may contribute to the impaired vascular morphology and function observed following TGF-β stimulation. GO enrichment analyses of the remaining DEG subsets are shown in Supplementary Fig. 5.

### In vivo vascular morphological changes by suppression of TGF-β signals

To investigate the role of TGF-β signaling in vascular morphogenesis *in vivo*, we performed quantitative morphological analyses of blood vessels in TβRII^iΔEC^ and their control, TβRII^fl/fl^. PECAM-1 fluorescence images revealed that the vascular network of TβRII^iΔEC^ mice exhibited a thickened vascular network (Figs. 4a and 4b). These findings demonstrate that the loss of endothelial TGF-β signaling leads to excessive vessel enlargement *in vivo*. Consistent with this regulatory role, TGF-β stimulation in our *in vitro* 3D vascular models led to a marked reduction in vascular area and diameter in both the 3D-Co and 3D-CM platforms (Figs. 1b, 1c, 2b, and 2c), while the inhibition of TGF-β signaling increased the vascular area and diameter (Figs. 4c and 4d). These findings support a model in which TGF-β signaling plays a critical role in maintaining vascular morphological homeostasis, with both gain- and loss-of-function perturbations leading to pronounced structural alterations. Our 3D vascular model recapitulates the morphological responses to TGF-β signaling observed *in vivo*, providing a quantitative *in vitro* platform for investigating TGF-β-mediated vascular remodeling.

**Fig. 4.**
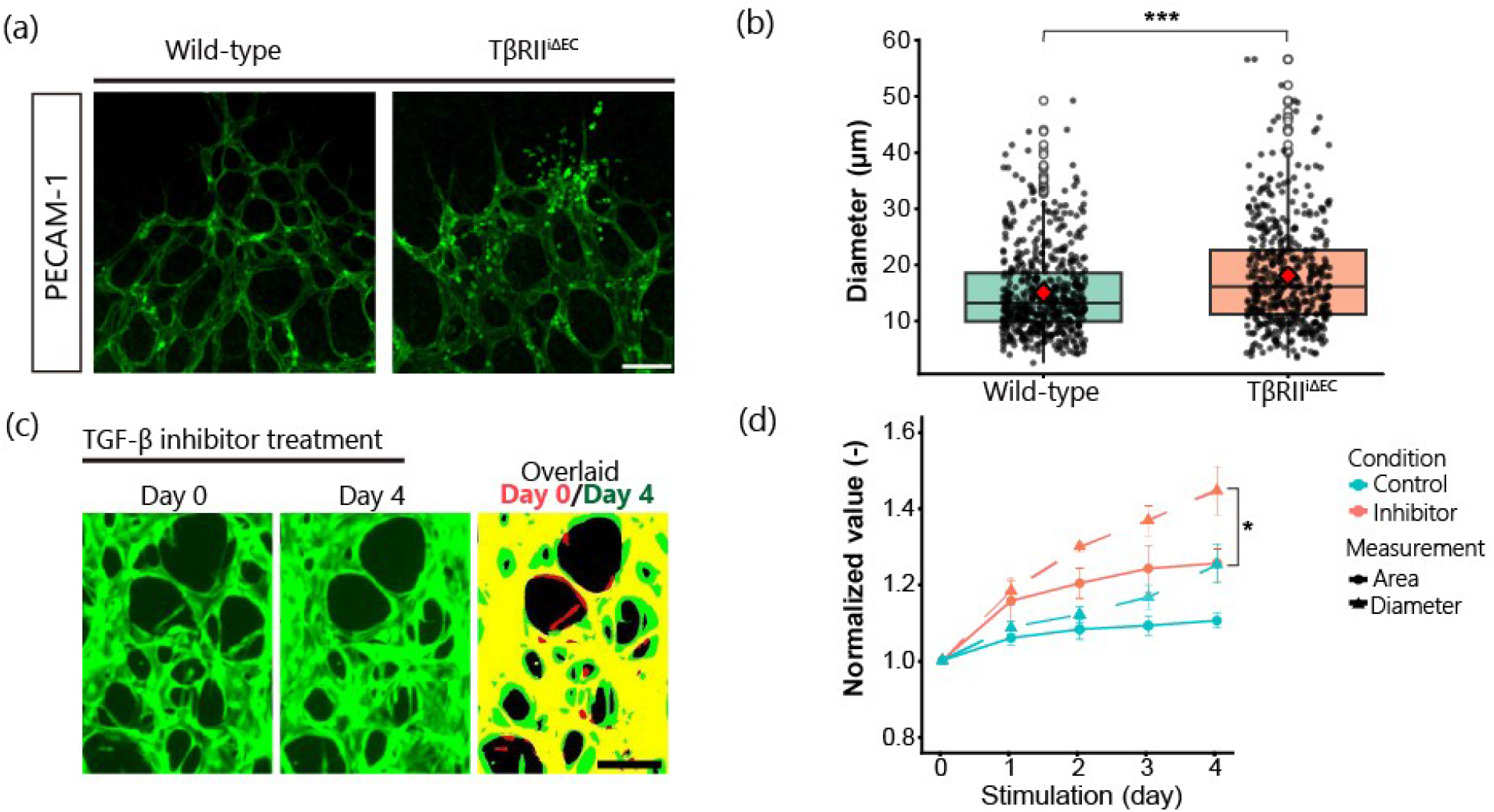
Vascular alterations between wild type and endothelial cell-specific TGF-β type II receptor-deficient mice (TβRII^iΔEC^) regulated by TGF-β *in vivo* and 3D-Co. (a) PECAM-1-stained (green) images of the anterior vasculature. Scale bar: 100 μm. (b) Vascular diameter distribution *in vivo*, n = 3. Statistical analyses: Mann–Whitney U test; \*\*\**p* < 0.001. (c) Representative images showing an elevation in the vascular area formed by GFP-HUVECs after treatment with TGF-β inhibitor in 3D-Co. In the overlaid image, red indicates the binarized vascular area at day 0 (start of stimulation), and green represents the binarized vascular area at day 4 (end of stimulation). Overlapping regions appear yellow. Scale bar: 200 μm. (d) Changes in vascular area and diameter after treatment with TGF-β inhibitor. All quantifications were normalized to the day 0 values. Statistical analyses: one-way ANOVA; \**p* < 0.05. Comparisons without asterisks are not statistically significant.

## Discussion

In this study, we utilized a 3D vascular model to investigate endothelial responses to TGF-β2 (e.g., EndoMT and vascular remodeling) to better recapitulate the *in vivo* microenvironment compared to conventional 2D systems. Our findings demonstrate that the 3D context is a critical determinant of endothelial responsiveness to TGF-β, enabling the emergence of structural and functional phenotypes that are not readily observed in 2D cultures. This highlights the importance of spatial architecture in studying complex vascular pathophysiology.

A prominent phenotype observed in this study was the reduction in vascular area and diameter following TGF-β stimulation. This structural regression was accompanied by increased apoptosis, enhanced EndoMT, and reduced endothelial proliferative activity, which is consistent with previous reports describing TGF-β-mediated inhibition of endothelial proliferation^34^, apoptosis induction^33^, and EndoMT promotion^4^. These findings suggest that TGF-β signaling contributes to vascular remodeling by shifting ECs toward a mesenchymal-like state, potentially leading to reduced structural stability and vascular thinning. However, whether vascular regression is directly driven by EndoMT or occurs through parallel mechanisms remains unclear.

Transcriptomic analysis provides further insights into how the 3D vascular microenvironment shapes endothelial cell behavior. Previous comparisons between 2D and 3D endothelial cultures have primarily focused on blood–brain barrier models, often involving co-culture with supporting cells, such as pericytes or astrocytes^35–37^. Consequently, it has remained difficult to distinguish the intrinsic effects of vascular architecture from those introduced by the surrounding cell types. In this study, the 3D-CM model enabled a direct comparison of ECs cultured under 2D and 3D conditions within the same biological system to isolate the influence of vascular architecture on endothelial transcriptional programs. Compared with 2D cultures, the 3D model displayed reduced basal expression of genes associated with TGF-β signaling, cell proliferation, and endothelial migration pathways, suggesting that ECs organized into a 3D vascular network adopted a more quiescent and metabolically distinct basal state than conventional 2D monolayers. The distinct responses observed in 3D vascular networks may result from vascular architecture-specific cues, including lumen formation, cell-cell interactions, and extracellular matrix engagement, which are absent in conventional 2D cultures. Such differences may arise from the reconstruction of capillary-like vascular architecture rather than differences in soluble microenvironmental cues, as all comparisons were performed within the same endothelial culture system. Following the TGF-β stimulation, gene expression profiling revealed coordinated upregulation of pathways associated with cell migration, extracellular matrix organization, inflammatory signaling, and apoptosis, along with the downregulation of cell cycle-related processes across both 2D and 3D cultures. These changes are consistent with the role of TGF-β in promoting endothelial remodeling and EndoMT. We observed a distinct metabolic signature in the 3D system characterized by reduced lipid biosynthesis and enhanced mitochondrial oxidative pathways. This may represent a previously unrecognized feature of TGF-β responses that becomes apparent only within a physiologically relevant 3D vascular architecture. However, because microfluidic devices contain only a limited volume of culture medium, nutrient availability and metabolite accumulation may differ substantially from conventional culture systems, potentially influencing cellular metabolic states. Therefore, the observed increase in mitochondrial oxidative pathways may reflect not only EndoMT or vascular remodeling but also adaptation to the unique microenvironment of the microfluidic culture system. Further studies are required to determine their causal contribution to the observed phenotypes. Besides, although HUVECs were used as a widely accepted endothelial model in this study, future studies using organ-specific endothelial cells will be valuable to further validate the generalizability of these findings.

Functionally, TGF-β stimulation led to a significant increase in vascular permeability in the 3D models. Given the essential role of endothelial junction integrity in maintaining barrier function, this finding suggests that TGF-β disrupts endothelial cohesion, potentially through cytoskeletal remodeling and junctional disassembly associated with EndoMT. Increased permeability is a hallmark of vascular dysfunction and is frequently associated with inflammatory conditions and pathological remodeling^38,39^. Therefore, the functional impairment observed in our model further supports the relevance of this system for studying disease-associated vascular changes.

Importantly, the vascular morphological changes observed in our 3D model were consistent with those identified in TβRII^iΔEC^ mice. Whereas the suppression of endothelial TGF-β signaling *in vivo* resulted in excessive vascular enlargement, the activation of TGF-β signaling in the 3D model induced vascular regression, together supporting a conserved role of TGF-β signaling in maintaining vascular morphological homeostasis. Although our study does not directly demonstrate that the 3D model fully reproduces the *in vivo* endothelial transcriptome, the ability to recapitulate key morphological features of TGF-β-dependent vascular remodeling highlights its potential as a physiologically relevant platform for investigating vascular disease mechanisms.

Our results suggest that the reconstruction of a self-organized 3D vascular architecture not only reproduces key morphological features of vascular remodeling and EndoMT but also establishes a distinct endothelial transcriptional state that cannot be captured by conventional 2D monolayer culture. These findings highlight the importance of vascular tissue architecture in regulating endothelial responses to TGF-β and support the application of vascularized microphysiological systems for mechanistic studies and therapeutic strategies targeting endothelial dysfunction.

## Conclusion

In this study, we utilized a physiologically relevant 3D vascular network platform to investigate TGF-β-induced EndoMT and vascular remodeling. Using this system, we demonstrated that TGF-β stimulation at an optimized concentration of 3 ng/mL induces pronounced structural regression and functional impairment of vascular networks, as evidenced by the increased SM22α expression, enhanced permeability, and network fragmentation.

Both 3D-Co and 3D-CM models consistently recapitulated key features of EndoMT. Transcriptomic profiling further revealed that TGF-β stimulation promotes coordinated changes in pathways associated with cell migration, extracellular matrix remodeling, inflammatory signaling, and apoptosis, accompanied by the suppression of cell cycle progression, collectively indicating a shift toward a mesenchymal-like and dysfunctional endothelial state.

Comparative analysis across culture dimensions revealed that ECs established fundamentally distinct basal transcriptional states in 2D and 3D environments even before TGF-β stimulation. Compared with conventional 2D cultures, the 3D vascular network exhibited a more quiescent basal phenotype with a distinct transcriptional response to TGF-β characterized by metabolic reprogramming, including reduced lipid biosynthesis and enhanced mitochondrial oxidative pathways, suggesting that spatial context critically shapes endothelial responsiveness to TGF-β signaling.

Our findings highlight the essential role of the 3D microenvironment in modulating TGF-β-driven vascular remodeling and provide a robust *in vitro* framework for dissecting the complex mechanisms underlying vascular pathology.

## Acknowledgments

This work was supported by the Japan Society for the Promotion of Science (JSPS) KAKENHI (Grant numbers 23H01821, 24K0261, 24K0203, 25H01351), JST FOREST Program (JPMJFR234S), JST SPRING (JPMJSP2180), AMED (JP24ama221238h0001), Tokyo Tech-TMDU Matching Fund, Asahi Glass Foundation, and MSD Life Science Foundation. We appreciate Ms. Maki Kamimura’s assistance in fabricating microfluidic devices and extracting total RNAs from microfluidic devices. This work was supported by the “Advanced Research Infrastructure for Materials and Nanotechnology in Japan (ARIM)” of the Ministry of Education, Culture, Sports, Science, and Technology (MEXT), Grant Numbers JPMXP1225UT1111, JPMXP1224UT1099, and JPMXP1223UT1172. All photolithography was conducted in the Takeda Super Cleanroom, Center of The University of Tokyo for The Advanced Research Infrastructure for Materials and Data Hub. The authors thank the Biomedical Center, Institute of Science Tokyo for the confocal microscopy analysis.

## Author contributions

Y.F.: Conceptualization, Methodology, Investigation, Formal analysis, Visualization, Writing – original draft. K. Tsuchiya: Methodology, Investigation, Formal analysis, Visualization, Writing – review & editing.

Y.N.: Conceptualization, Methodology, Formal analysis, Supervision, Funding acquisition, Writing – review & editing. K. Takahashi: Formal analysis, Interpretation of data, Writing – review & editing.

Y.O.: Investigation, Writing – review & editing.

S.K.: Investigation, Writing– review & editing. S.Y.: Investigation, Writing – review & editing. M.K.: Investigation, review & editing.

F.I.: Resources, Writing – review & editing.

T.W.: Conceptualization, Supervision, Writing – review & editing.

H.K.: Conceptualization, Funding acquisition, Project administration, Supervision, Writing – review & editing.

## References

1. Xiao, Y. & Yu, D. Tumor microenvironment as a therapeutic target in cancer. Pharmacol Ther 221, 107753 (2021).

2. Fukumura, D., Kloepper, J., Amoozgar, Z., Duda, D. G. & Jain, R. K. Enhancing cancer immunotherapy using antiangiogenics: opportunities and challenges. Nat Rev Clin Oncol 15, 325–340 (2018).

3. Ma, J., Sanchez-Duffhues, G., Goumans, M.-J. & ten Dijke, P. TGF-β-Induced Endothelial to Mesenchymal Transition in Disease and Tissue Engineering. Front. Cell Dev. Biol. 8, (2020).

4. Watabe, T., Takahashi, K., Pietras, K. & Yoshimatsu, Y. Roles of TGF-β signals in tumor microenvironment via regulation of the formation and plasticity of vascular system. Semin Cancer Biol 92, 130–138 (2023).

5. Zeisberg, E. M. et al. Endothelial-to-mesenchymal transition contributes to cardiac fibrosis. Nat. Med. 13, 952–961 (2007).

6. Akatsu, Y. et al. Fibroblast growth factor signals regulate transforming growth factor-β-induced endothelial-to-myofibroblast transition of tumor endothelial cells via Elk1. Molecular Oncology 13, 1706–1724 (2019).

7. Katsura, A. et al. MicroRNA-31 is a positive modulator of endothelial–mesenchymal transition and associated secretory phenotype induced by TGF-β. Genes to Cells 21, 99–116 (2016).

8. Kokudo, T. et al. Snail is required for TGFβ-induced endothelial-mesenchymal transition of embryonic stem cell-derived endothelial cells. Journal of Cell Science 121, 3317–3324 (2008).

9. TGF-β-induced mesenchymal transition of MS-1…: Journal of Biochemistry. https://www.ovid.com/journals/jbioc/abstract/10.1093/jb/mvr121~tgfinduced-mesenchymal-transition-of-ms1-endothelial-cells.

10. Suzuki, H. I. et al. Regulation of TGF-β-mediated endothelial-mesenchymal transition by microRNA-27. The Journal of Biochemistry 161, 417–420 (2017).

11. Takahashi, K. et al. CD40 is expressed in the subsets of endothelial cells undergoing partial endothelial– mesenchymal transition in tumor microenvironment. Cancer Science 115, 490–506 (2024).

12. Follain, G. et al. Hemodynamic Forces Tune the Arrest, Adhesion, and Extravasation of Circulating Tumor Cells. Dev Cell 45, 33–52.e12 (2018).

13. Winkler, J., Abisoye-Ogunniyan, A., Metcalf, K. J. & Werb, Z. Concepts of extracellular matrix remodelling in tumour progression and metastasis. Nat Commun 11, 5120 (2020).

14. Mohammadi, H. & Sahai, E. Mechanisms and impact of altered tumour mechanics. Nat Cell Biol 20, 766–774 (2018).

15. Ingber, D. E. Human organs-on-chips for disease modelling, drug development and personalized medicine. Nat Rev Genet 23, 467–491 (2022).

16. Reimagining human-centric drug development with new approach methodologies. https://www.science.org/doi/10.1126/science.aeb0045doi:10.1126/science.aeb0045.

17. Chandran Latha, K., et al. Shear Stress Alterations Activate BMP4/pSMAD5 Signaling and Induce Endothelial Mesenchymal Transition in Varicose Veins. Cells 10, 3563 (2021).

18. Karthika, C. L. et al. Oscillatory shear stress modulates Notch-mediated endothelial mesenchymal plasticity in cerebral arteriovenous malformations. Cell Mol Biol Lett 28, 22 (2023).

19. Mina, S. G. et al. Shear stress magnitude and transforming growth factor-βeta 1 regulate endothelial to mesenchymal transformation in a three-dimensional culture microfluidic device. RSC Adv. 6, 85457– 85467 (2016).

20. Mina, S. G., Huang, P., Murray, B. T. & Mahler, G. J. The role of shear stress and altered tissue properties on endothelial to mesenchymal transformation and tumor-endothelial cell interaction. Biomicrofluidics 11, 044104 (2017).

21. Breuil, L. et al. Vascular microphysiological systems (MPS): biologically relevant and potent models. Lab Chip 25, 4221–4251 (2025).

22. Offeddu, G. S. et al. An on-chip model of protein paracellular and transcellular permeability in the microcirculation. Biomaterials 212, 115–125 (2019).

23. Yeon, J. H. et al. Cancer-derived exosomes trigger endothelial to mesenchymal transition followed by the induction of cancer-associated fibroblasts. Acta Biomaterialia 76, 146–153 (2018).

24. Kim, S., Lee, H., Chung, M. & Jeon, N. L. Engineering of functional, perfusable 3D microvascular networks on a chip. Lab Chip 13, 1489 (2013).

25. Hanada, K. et al. Reduced lung metastasis in endothelial cell-specific transforming growth factor β type II receptor-deficient mice with decreased CD44 expression. iScience 27, 111502 (2024).

26. Schindelin, J., et al. Fiji: an open-source platform for biological-image analysis. Nat Methods 9, 676–682 (2012).

27. Lee, H., Kim, S., Chung, M., Kim, J. H. & Jeon, N. L. A bioengineered array of 3D microvessels for vascular permeability assay. Microvascular Research 91, 90–98 (2014).

28. Nashimoto, Y. et al. Integrating perfusable vascular networks with a three-dimensional tissue in a microfluidic device. Integrative Biology 9, 506–518 (2017).

29. Sabbineni, H., Verma, A. & Somanath, P. R. Isoform-specific effects of transforming growth factor β on endothelial-to-mesenchymal transition. J Cell Physiol 233, 8418–8428 (2018).

30. Derada Troletti, C., et al. Inflammation-induced endothelial to mesenchymal transition promotes brain endothelial cell dysfunction and occurs during multiple sclerosis pathophysiology. Cell Death Dis 10, 45 (2019).

31. Kovacic, J. C. et al. Endothelial to Mesenchymal Transition in Cardiovascular Disease. Journal of the American College of Cardiology 73, 190–209 (2019).

32. Piera-Velazquez, S., Li, Z. & Jimenez, S. A. Role of Endothelial-Mesenchymal Transition (EndoMT) in the Pathogenesis of Fibrotic Disorders. Am J Pathol 179, 1074–1080 (2011).

33. Van Meeteren, L. A. & Ten Dijke, P. Regulation of endothelial cell plasticity by TGF-β. Cell Tissue Res 347, 177–186 (2012).

34. Transforming growth factor-ß2 inhibition of corneal endothelial proliferation mediated by prostaglandin: Current Eye Research: Vol 26, No 6. https://www.tandfonline.com/doi/abs/10.1076/ceyr.26.5.363.15442.

35. Linville, R. M. et al. Three-dimensional microenvironment regulates gene expression, function, and tight junction dynamics of iPSC-derived blood–brain barrier microvessels. Fluids Barriers CNS 19, 87 (2022).

36. Sonninen, T.-M. et al. From inserts to chips: microfluidic culture and 3D astrocyte co-culture drive functional and transcriptomic changes in hiPSC-derived endothelial cells. Fluids Barriers CNS 22, 58 (2025).

37. Zhang, J. et al. A Genome-wide Analysis of Human Pluripotent Stem Cell-Derived Endothelial Cells in 2D or 3D Culture. Stem Cell Reports 8, 907–918 (2017).

38. Wilhelm, D. L. THE MEDIATION OF INCREASED VASCULAR PERMEABILITY IN INFLAMMATION. Pharmacological Reviews 14, 251–280 (1962).

39. Claesson-Welsh, L. Vascular permeability—the essentials. Upsala Journal of Medical Sciences 120, 135–143 (2015).

