## Supplementary File for "Three-dimensional vascular microenvironments uncover endothelial plasticity during TGF-β2-driven vascular remodeling"

**Supporting Information**

Contents

1. Methods

2. Figures

*Formation of a 3D vascular network and corresponding 2D cell culture*

To create a 3D co-culture vascular network (3D-Co), GFP-HUVECs and hLFs were seeded into the microfluidic device. The microfluidic device comprises five parallel channels separated by trapezoidal or hexagonal microposts, facilitating the precise patterning of hydrogels. GFP-HUVECs and hLFs were suspended in bovine fibrinogen solution (Merck, final concentration: 3.0 mg/mL) at $8.0 \times 10^6$ and $5.0 \times 10^6$ cells/mL, respectively. After mixing with bovine thrombin (Merck, final concentration: 0.5 U/mL), GFP-HUVECs were introduced to Channel 3 (Ch. 3), and hLFs were introduced to Chs. 1 and 5 (Fig. 1a). The microfluidic device was then incubated at 37 °C for 15 min to enhance the polymerization of fibrinogen and to form fibrin gel to support GFP-HUVECs and hLFs. After incubation, EGM-2 was introduced into Chs. 2 and 4. Half of the medium in Chs. 2 and 4 were changed daily. To investigate the permeability coefficient of the vascular network and enhance the connection between the microchannels and vascular network, additional GFP-HUVECs were introduced into Chs. 2 and 4 and attached to both sides of the fibrin gel in Ch. 3 on the second day after cell introduction into the device. After seven days in the device culture, the 3D-Co was formed in a microfluidic device and used for TGF-β stimulation.

To clarify the TGF-β effects on the 3D vascular network, excluding the effects of hLFs, we utilized the 3D vascular network formed by hLF conditioned media (CM). This model is hereafter designated as the 3D CM-driven vascular network (3D-CM). The confluent hLFs in a 100-mm culture dish were prepared to collect the CM. Two mL of EGM-2 was added to the hLF dish, and one day after incubation, the CM was collected and used for the device culture immediately. hLFs in a 100-mm culture dish were continuously used to collect the CM daily for more than 10 days. GFP-HUVECs embedded in fibrin gel were introduced into Ch. 3, similar to the vascular formation with hLFs (see above), and the blank fibrin gel was filled with Chs. 1 and 5. The collected CM was introduced to Chs. 2 and 4 (Fig. 2a). Half of the medium in Chs. 2 and 4 were exchanged daily.

TGF-β2 (hereafter termed TGF-β) treatment was performed by adding TGF-β2 (Thermo Fisher Scientific) to EGM-2 and incubating for more than 72 h. The final concentration of TGF-β was 0, 1, 3 and 10 ng/mL. As the buffer for TGF-β, 0.1% BSA in 4 mM HCl was added at 0.1% of the culture medium volume. The same volume was also added to the control sample (0 ng/mL TGF-β). TGF-β was used at concentrations of 0, 1, 3, and 10 ng/mL in 3D-Co, while concentrations of 0 and 3 ng/mL were used in the 3D-CM.

To compare the 3D-CM model with the 2D culture, GFP-HUVECs were seeded into 35 mm dishes at a density of $1.6 \times 10^5$ cells/dish. Because cell-cell junctions may influence the development of EndoMT, both confluent (100% confluence) and sparse (60%–70% confluence) samples were prepared. Sparse samples were stimulated starting on day 2 after seeding (day 0), while confluent samples were stimulated on day 4. TGF-β was applied at concentrations of 0 and 3 ng/mL for 72 h. Filtered CM containing TGF-β was replaced every other day throughout the stimulation period.

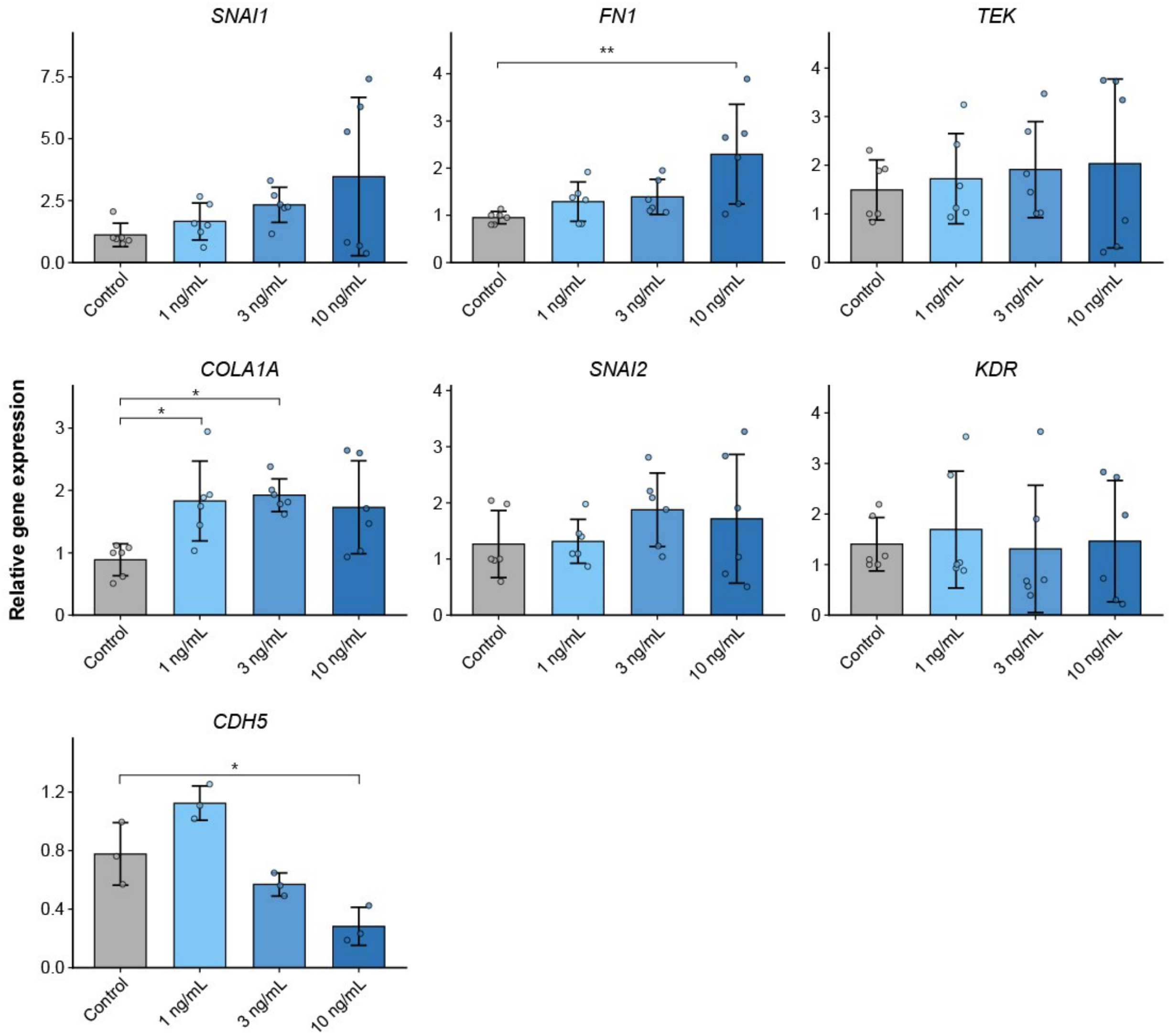


Supplementary Fig. 1 RT-qPCR analysis of gene expression in the 3D vascular network under TGF-β stimulation. Gene expression levels were quantified in 3D-Co treated with TGF-β2 (1, 3, and 10 ng/mL) for four days, with untreated samples serving as the control (0 ng/mL). The panels show the expression of mesenchymal markers (e.g., *SNAI1*, *SNAI2*, *FN1*, and *COL1A1*) and endothelial markers (e.g., *KDR*, *CDH5*, and *TEK*). Data are presented as mean ± standard deviation. Statistical significance was determined using one-way ANOVA followed by Tukey–Kramer HSD post-hoc test. *$p < 0.05$, **$p < 0.01$ versus the control group. Comparisons without asterisks are not statistically significant.

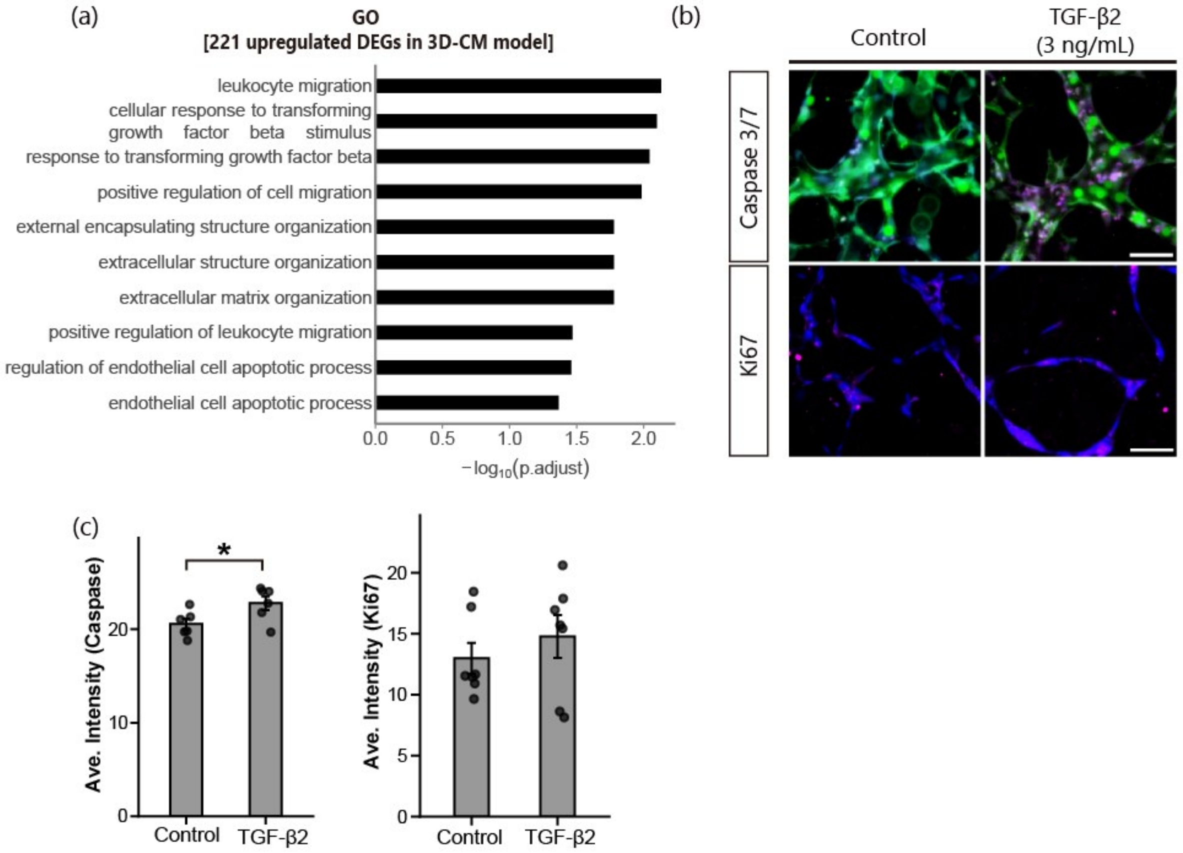


Supplementary Fig. 2 (a) Gene ontology functional enrichment analysis of differentially expressed genes upregulated by TGF-β in the 3D-CM model. (b) Immunocytochemical analysis. Staining for caspase 3/7 (magenta) and nuclei (cyan), Ki67 (magenta) and nuclei (blue). Green is GFP. Scale bar: 100 μm. (c) Average intensity of caspase 3/7 (left) and Ki67 (right). Data are presented as mean ± standard deviation; Statistical significance was determined by Student's t-test; $*p < 0.05$; n = 6 independent samples. Comparisons without asterisks are not statistically significant.

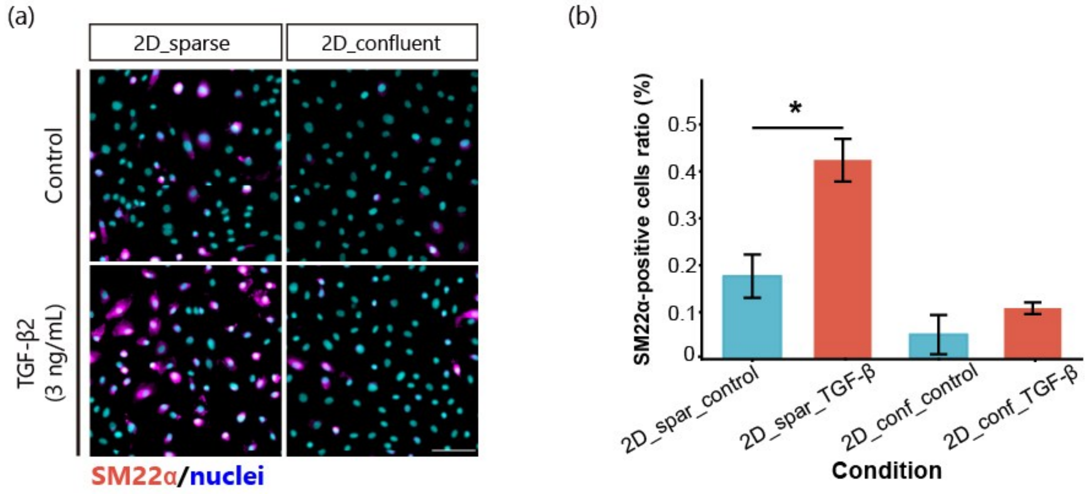


Supplementary Fig. 3 Immunocytochemical analysis of GPF-HUVECs in 2D sparse and confluent culture. (a) Staining for SM22α (magenta) and nuclei (cyan). Scale bar: 100 µm. (b) Average intensity of SM22α, n = 6. All data are shown as mean ± standard deviation. Statistical analyses: t-test; $*p < 0.05$. Comparisons without asterisks are not statistically significant.

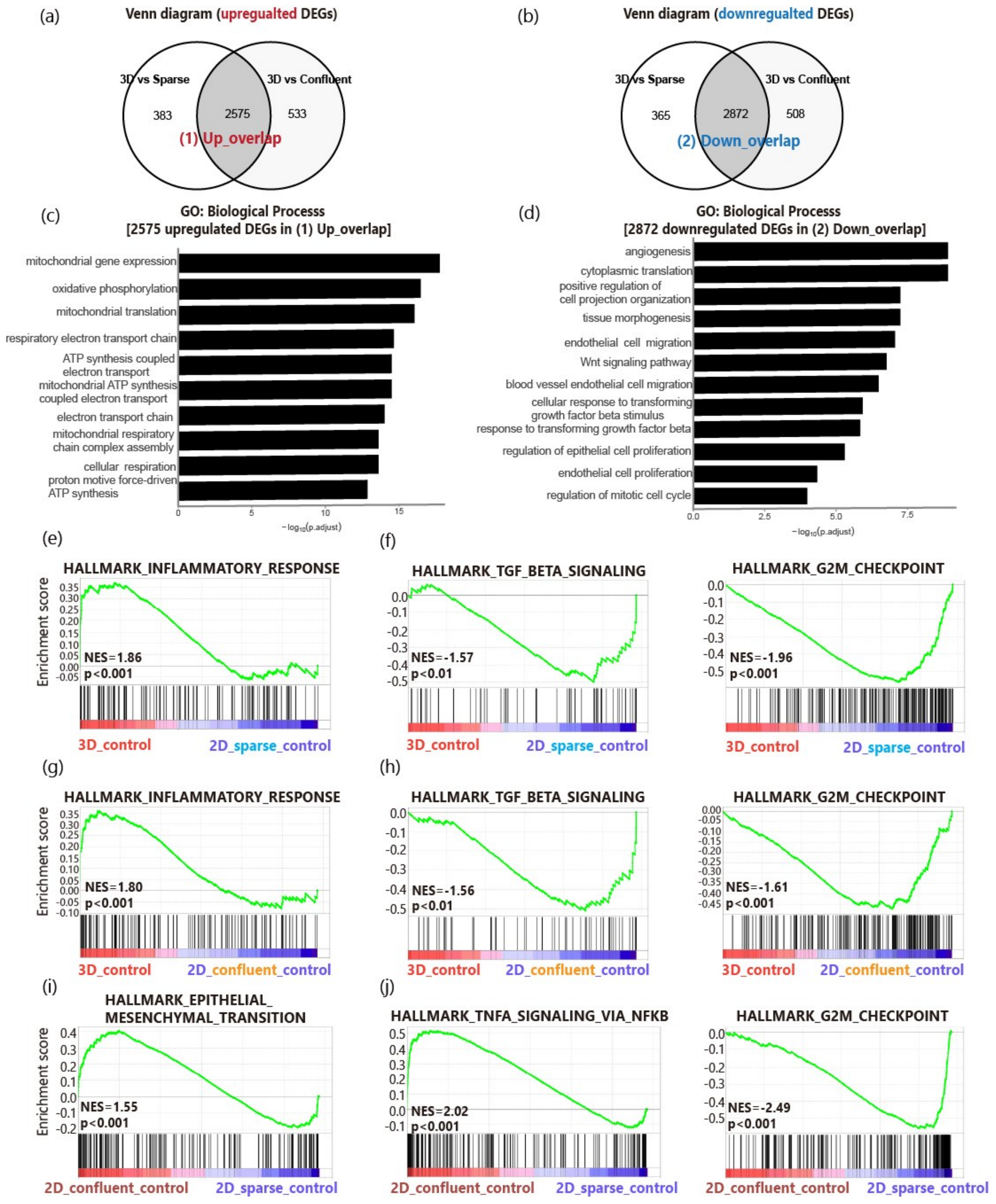


Supplementary Fig. 4 Comparative RNA-seq analysis of nonstimulated control groups in 2D sparse, 2D confluent, and 3D-CM culture models. Venn diagrams showing shared (a) upregulated and (b) downregulated differentially expressed genes (DEGs) identified from comparisons of 3D control vs. 2D sparse control and 3D control vs. 2D confluent control. The overlapping gene sets were designated as (1) Up_overlap and (2) Down_overlap. Gene ontology enrichment analysis of the (c) 2,575 DEGs in Up_overlap and (d) 2,872 DEGs in Down_overlap, with the top 10 enriched biological processes ranked by adjusted *p* value. Gene set enrichment analysis comparing (e, f) 3D control with 2D sparse control, (g, h) 3D control with 2D confluent control, and (i, j) 2D confluent control with 2D sparse control. n = 2 for the 3D group and n = 4 for the 2D groups. NES, normalized enrichment score; *q*, false discovery rate-adjusted *p* value.

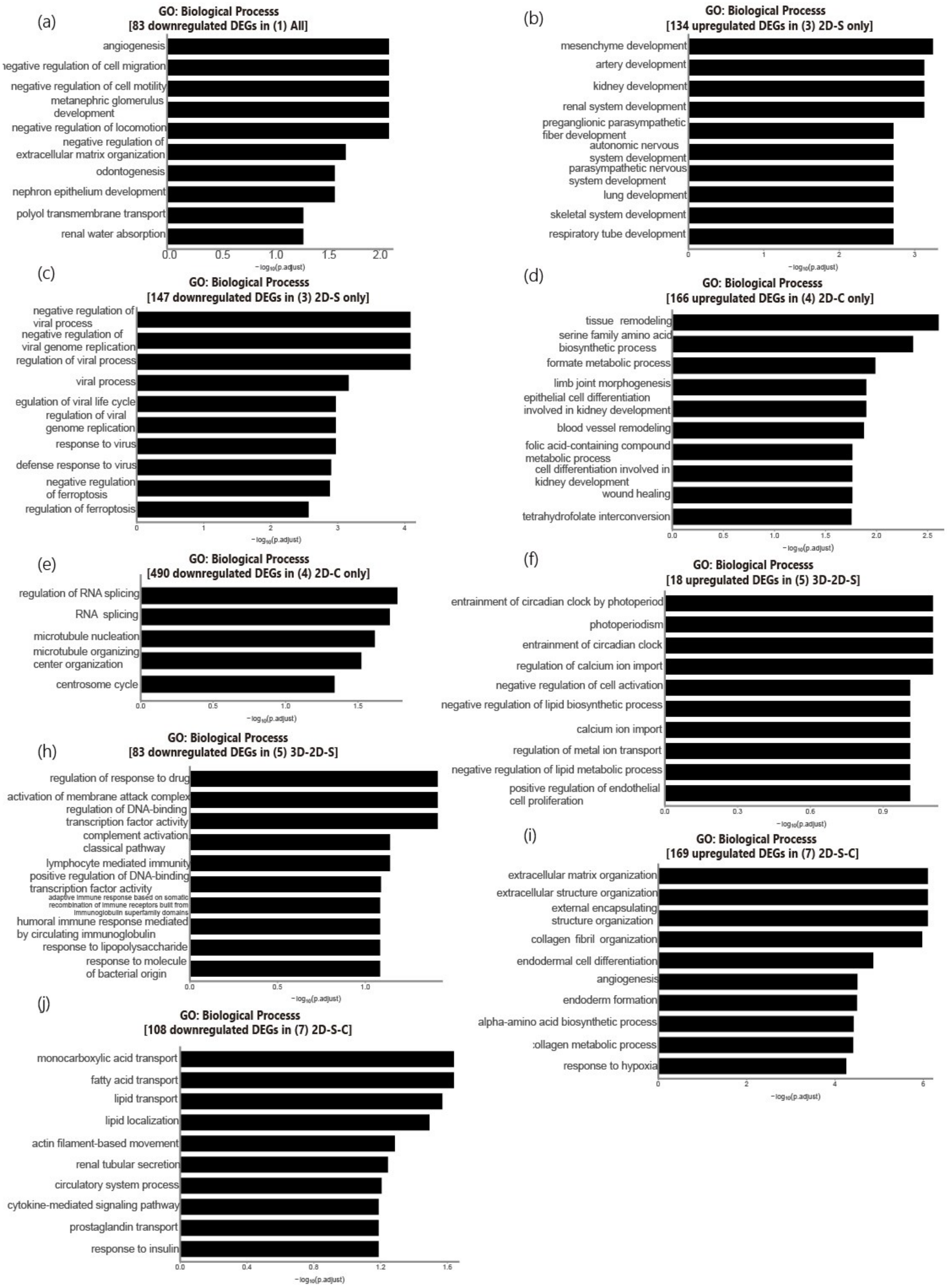

(a)
GO: Biological Processs
[83 downregulated DEGs in (1) All]
angiogenesis
egative regulation of cell migration
negative regulation of cell motility
metanephric glomerulus development
negative regulation of locomotion
negative regulation of extracellular matrix organization
odontogenesis
nephron epithelium development
polyol transmembrane transport
renal water absorption
0.0 0.5 1.0 1.5 2.0
−log10(p.adjust)
(b)
GO: Biological Processs
[134 upregulated DEGs in (3) 2D-S only]
mesenchyme development
artery development
kidney development
renal system development
preganglionic parasympathetic fiber development
autonomic nervous system development
parasympathetic nervous system development
lung development
skeletal system development
respiratory tube development
0 1 2 3
−log10(p.adjust)
(c)
GO: Biological Processs
[147 downregulated DEGs in (3) 2D-S only]
negative regulation of viral process
negative regulation of viral genome replication
regulation of viral process
viral process
egulation of viral life cycle
regulation of viral genome replication
response to virus
defense response to virus
negative regulation of ferroptosis
regulation of ferroptosis
0 1 2 3 4
−log10(p.adjust)
(d)
GO: Biological Processs
[166 upregulated DEGs in (4) 2D-C only]
tissue remodeling
serine family amino acid biosynthetic process
formate metabolic process
limb joint morphogenesis
epithelial cell differentiation involved in kidney development
blood vessel remodeling
folic acid-containing compound metabolic process
cell differentiation involved in kidney development
wound healing
tetrahydrofolate interconversion
0.0 0.5 1.0 1.5 2.0 2.5
−log10(p.adjust)
(e)
GO: Biological Processs
[490 downregulated DEGs in (4) 2D-C only]
regulation of RNA splicing
RNA splicing
microtubule nucleation
microtubule organizing center organization
centrosome cycle
0.0 0.5 1.0 1.5
−log10(p.adjust)
(f)
GO: Biological Processs
[18 upregulated DEGs in (5) 3D-2D-S]
entrainment of circadian clock by photoperiod
photoperiodism
entrainment of circadian clock
regulation of calcium ion import
negative regulation of cell activation
negative regulation of lipid biosynthetic process
calcium ion import
regulation of metal ion transport
negative regulation of lipid metabolic process
positive regulation of endothelial cell proliferation
0.0 0.3 0.6 0.9
−log10(p.adjust)
(h)
GO: Biological Processs
[83 downregulated DEGs in (5) 3D-2D-S]
regulation of response to drug
activation of membrane attack complex
regulation of DNA-binding transcription factor activity
complement activation classical pathway
lymphocyte mediated immunity
positive regulation of DNA-binding transcription factor activity
adaptive immune response based on somatic recombination of immune receptors built from immunoglobulin superfamily domains
humoral immune response mediated by circulating immunoglobulin
response to lipopolysaccharide
response to molecule of bacterial origin
0.0 0.5 1.0
−log10(p.adjust)
(i)
GO: Biological Processs
[169 upregulated DEGs in (7) 2D-S-C]
extracellular matrix organization
extracellular structure organization
external encapsulating structure organization
collagen fibril organization
endodermal cell differentiation
angiogenesis
endoderm formation
alpha-amino acid biosynthetic process
ollagen metabolic process
response to hypoxia
0 2 4 6
−log10(p.adjust)
(j)
GO: Biological Processs
[108 downregulated DEGs in (7) 2D-S-C]
monocarboxylic acid transport
fatty acid transport
lipid transport
lipid localization
actin filament-based movement
renal tubular secretion
circulatory system process
cytokine-mediated signaling pathway
prostaglandin transport
response to insulin
0.0 0.4 0.8 1.2 1.6
−log10(p.adjust)

Supplementary Fig. 5 Gene ontology (GO) enrichment analysis of differentially expressed genes from each region of the Venn diagrams shown in Fig. 3c and d. Panels (a–j) present the top 10 enriched biological processes ranked by adjusted *p* value for each gene set. No significant GO enrichment was identified for gene set (6) (3D–2D-C).

Supplementary Table 1. Primer sequences used for quantitative real-time PCR.

| Gene | Forward (5′–3′) | Reverse (5′–3′) |
|---|---|---|
| *TAGLN* | TCAAGCAGATGGAGCAGGTG | GCTGCCATGTCTTTGCCTTC |
| *COL1A1* | GCTCTTGCAACATCTCCCCT | CCTTCCTGACTCTCCTCCGA |
| *FN1* | AAACCAATTCTTGGAGCAGG | CCATAAAGGGCAACCAAGAG |
| *SNAI1* | TTCTCACTGCCATGGAATTCC | GCAGAGGACACAGAACCAGAAA |
| *SNAI2* | GCCTCCAAAAAGCCAAACTACA | GAGGATCTCTGGTTGTGGTATGACA |
| *TEK* | GGTGGAAAAGCCCTTCAACA | CATCCCCAAAGTAAGGCTCAG |
| *KDR* | CAGAATCCCTGCGAAGTACCTT | GTCAGTACATGCCCCGCTTTAA |
| *CLDN5* | CTCCCCAGGCTTATCCAACG | CGGCGACTACGACAAGAAGA |
| *CDH5* | TAGCATTGGATACTCCATCCGC | GCCGTGTTATCGTGATTATCCG |
| *β-Actin* | TCACCCACACTGTGCCCATCTACA | CAGCGGAACCGCTCATTGCCAATGG |
